# Branching responses to pruning in cocoa

**DOI:** 10.1101/2023.03.08.531700

**Authors:** Ambra Tosto, Jochem B. Evers, Niels P. R. Anten, Pieter A. Zuidema

## Abstract

The branching pattern of a tree determines the efficiency of light interception and carbon assimilation. Pruning can modify the branching pattern, as a result of changes in physiological and environmental conditions, and ultimately pruning can have major effects on yield. For one of the major tropical tree crops, cocoa (*Theobroma cacao*), very little is known about branching response to pruning. To address this knowledge gap we performed a pruning experiment on young cocoa trees in Côte d’Ivoire.

We applied five treatments: two heading treatments (the removal of the terminal apex or 66% of a branch) and two thinning treatments (the removal of 1 or 2 primary branches) and one unpruned control. The branching pattern of the primary branches was described by the number, position, and length of lateral branches right after pruning, and the same observations were repeated after a cycle of leaf production. The probability of branching and the length of lateral branches along a primary branch, in pruned and unpruned conditions, was analyzed using generalized mixed effect models.

In unpruned conditions, the probability of branch presence was higher towards the middle of the primary branches and lower at the extremes. Branch length decreased going from the base to the tip of a primary branch. After one cycle of leaf production, new branches emerged preferentially on the distal section of a branch, but probability of branch emergence was reduced by the presence of other lateral branches. Pruning increased the probability of branch emergence mostly towards the tip of a branch, with heavy heading having the strongest effect. By contrast, heavy thinning increased branch emergence also toward the base of the branch.

Our results can be applied to improve formation pruning, as this may trigger branching in different part of the crown, depending on the form of pruning. Our study also assists the development of three-dimensional tree models that could further our understanding of the impact of pruning on cocoa growth and productivity.

## 1. Introduction

One of the most important determinants of tree architecture is the branching pattern of the tree. The appearance, location, and size of the branches determine the distribution of the foliage, influencing the efficiency of light interception and thus whole-plant photosynthesis (Niinemets, 2010). The branching pattern is the result of the interaction between genes, environmental conditions and perturbations (McSteen & Leyser, 2005a; Sachs & Novoplansky, 1995). Interspecific differences in branching patterns also reflect divergent adaptations to different habitats (Poorter et al., 2006).

Branching is regulated by correlative inhibition. This comprises various forms of suppressive signaling among organs. One of the best-known forms of correlative inhibition is apical dominance whereby an apical bud exerts a suppressive signal on lateral buds in basipetal direction. The presence of other organs also contributes to the inhibition of bud break and branch growth. This inhibition can be exerted by organs on the same branch (within-branch inhibition) or by nearby branches (between-branch correlative inhibition) (Wubs et al., 2013; Zieslin & Halevy, 1976).

Pruning is an agricultural or horticultural practice mostly used to modify the architecture of woody perennials, such as tree crops and ornamental trees, to control tree size, increase light capture efficiency, increase yield and/or fruit quality, facilitate harvesting and spraying activities or for esthetic purposes (Ferree & Schupp, 2003). Pruning interventions typically consist of a combination of heading and thinning cuts. A heading cut removes a portion of a branch including the apex. This releases the apical dominance stimulating bud outgrowth and branch vigor in the remaining branch section (Wilson, 2000). In contrast, a thinning cut removes an entire branch, reducing between-branch correlative inhibition. This can result in the outgrowth of some buds on the remaining branches, but compared to a heading cut, the response triggered by a thinning cut is generally found to be weaker (Ferree & Schupp, 2003).

Pruning induced branching and increased branch vigor are part of the suite of compensatory mechanisms with which trees mitigate the negative effect of biomass removal (Anten et al., 2003). Up to a certain level, compensatory responses become stronger as biomass removal increases. For instance, in apple and mango trees the length of the new branches increases with the intensity of the heading cut (Mika, 1986, Persello et al., 2019).

The influence of pruning on branching has been studied in several perennial crops in temperate (Fumey et al., 2011; Li et al., 2010; Marini et al., 2020; Wubs et al., 2013) and tropical climates (Persello et al., 2019). However, to date, very limited information on branching pattern and branching responses to pruning is available for cocoa (*Theobroma cacao*), one of the most important tropical tree crops. (Asante et al., 2022; Fairtrade Foundation, 2016). Pruning of cocoa is considered an important yield-enhancing practice, while it may also help with tree management (e.g., disease control) and, as such, is recommended to farmers. Two main types of pruning are performed in cocoa: formation pruning to establish the structure of the crown after the tree has developed the first whorl of branches; and maintenance pruning, that is performed one or more times per year throughout the tree life-cycle, to reduce excessive self-shading and constrain tree dimensions (IITA, 2020). Scientific evidence on the effect of formation pruning is scarce (KAU, 1988, 1989, 1991) and evidence of the effect of maintenance pruning on yield is mixed (Tosto et al., 2022a). Additionally, pruning recommendations tend to be very general. This has resulted in low adoption of the practice (Obeng Adomaa et al., 2022). Therefore, understanding the branching response of cocoa to pruning may contribute to the development of effective, yield-increasing cocoa pruning practices.

We addressed the following questions: (1) To what extent do heading cuts affect the probability of branching? (2) To what extent is branching probability affected by thinning cuts? (3) To what extent does the type and intensity of pruning affect biomass allocation to lateral branches?

To answer these questions we first analyzed branching pattern and biomass allocation in unpruned trees. Since cocoa is originally a shade-adapted species (Lachenaud et al., 2005), we hypothesize that cocoa will exhibit a branching pattern that minimizes leaf overlap. We expect therefore the trees to have more and longer branches in the distal sections of the primary branches compared to the more proximal sections. When pruning is applied, we hypothesize that heading cuts (question 1) will strongly stimulate branching, especially in the distal sections of a primary branch due to the removal of apical dominance. Additionally, we hypothesize that thinning cuts (question 2) will induce a weak branching response, mostly concentrated at the base of the remaining branching due to a decrease in correlative inhibition. Finally, we hypothesize that more intensive pruning (i.e., removing more biomass, question 3) will result in increased allocated to new and existing lateral branches as a result of compensatory growth mechanisms.

To test our hypotheses, we applied two levels of heading and thinning cuts to the primary plagiotropic branches of two-year-old field-grown cocoa trees in an experimental plantation in Côte d’Ivoire. The branching pattern of pruned cocoa trees was quantified and compared with unpruned trees, and related to the intensity and type of pruning.

## 2. Methods

### 2.1. Study species

Cocoa is originally a shade-adapted understory species of the Amazon forest (Lachenaud et al., 2005). Its architectural development consists in the determinate growth of a vertical (orthotropic) shoot, at which end the young tree develops a whorl of horizontally spreading (plagiotropic) branches that form the so-called “jorquette” (Hallé et al., 1978). Under natural conditions, vertical growth consists in a buildup of several jorquettes, but in cultivated conditions, vertical growth is limited to one or two jorquette layers (Niemenak et al., 2009). In this study we only allowed for the developed of the first jorquette layer.

### 2.2. Study sites

This study was carried out at a research center located in the municipality of Tiassalé (5.913338 N, 4.867181 W), Côte d’Ivoire. Throughout the study period (2018-2021) the average daily temperature ranged from 22.1 and 31 °C and the mean annual cumulative precipitation was 1431 mm, with a dry season (<100 mm of precipitation per month) spanning approximately from December to March. Climate data were obtained from a weather station located ≈ 1 km from the study site. The soil there has a clay-loam texture and a pH at planting of 5.4.

### 2.3. Field planting design and field maintenance

Two adjacent parcels, A (0.41 ha) and B (1.08 ha), were established in a former young rubber plantation. In December 2017, we planted plantains (*Musa* sp.) and framire (*Terminalia ivorensis)*, a fast-growing tree species commonly used as shade trees in cocoa agroforestry systems, to create sufficient shade for the establishment of cocoa seedlings (see Figure S 1 for planting design). In June 2018, six-month-old cocoa seedlings were planted at a distance of 3 m by 3m (density of 1111 plants per ha, Figure S 1). In parcel A, a mix of Upper Amazonian hybrids was used, and in parcel B cocoa hybrids F1 and F2 (from *Centre National de Recherche Agronomique*, CNRA). Initially, only parcel A was intended to be used for the current study. However, due to plant mortality and slow development, too few suitable plants remained in parcel A in 2020. To increase sample size we therefore choose to include F1 plants from parcel B. In statistical analyses, parcel identity is explicitly accounted for.

At the moment of planting, each cocoa seedling received 50 g of Triple superphosphate, 150 g of N-P-K (15-15 -15) fertilizer and 10 liter of organic matter (mix of well-decomposed sawdust, rice husks, and chicken dung). Fertilization continued during the course of the experiment at a rate of 100 g/tree/year of N-P-K (15-15-15) applied twice a year (in May and in Oct).

Both parcels were equipped with an irrigation system with a dripping line along each cocoa-plantain-framire row. Supplementary irrigation was provided daily during establishment phase (June 2018-August 2018) and in the following years during drier months (December-March) to avoid water stress. Regular maintenance of the parcels (weeding, pesticide application) was carried out during the full duration of the experiment in both parcels. All suckers developing on the orthotropic stems of cocoa plants were removed, once per month. Once per year, lateral suckers of plantains were removed leaving only two pseudostems per plant to maintain shade to a stable level. In parcel B, shade levels were monitored throughout the study period with three Onset HOBO MX2202 light sensors (placed half a meter from a cocoa seedling). A reference sensor was placed in the proximity of the field in full sun. During the study period, the plantain-framire layer provided an average shade level of about 25%.

### 2.4. Treatments description and allocation

Our experiment had one control (unpruned) and four pruning treatments: two heading treatments and two thinning treatments. Heading treatments entailed the removal of either the apical bud (Head_tip) or 2/3 of the internodes (Head_66%) from all primary branches. Thinning treatments consisted of the complete removal of one (Thin_1) or two (Thin_2) primary branches. Control plants (Control) were left unpruned (figure 1).

**Figure 1.**
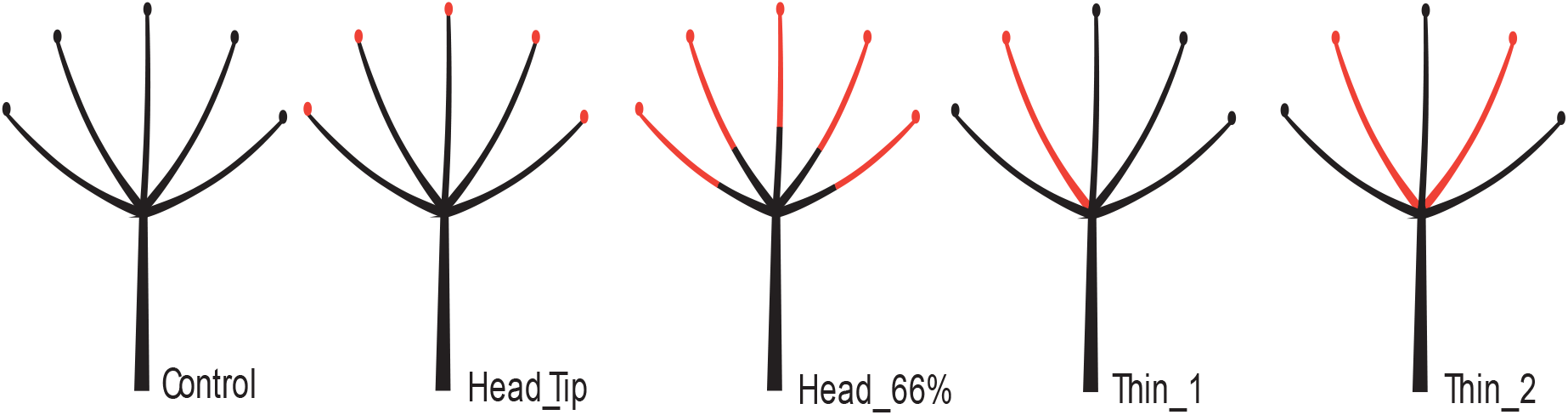
Schematic representation of pruning treatments. In red are sections removed by the treatments. Heading cuts: the removal of terminal apex (Head_tip); removal of branch section corresponding 66% of buds (Head_66%). Thinning cuts: removal of one primary branch (Thin_1); removal of two primary branches (Thin_2). Control plants were left unpruned.

Treatments were randomly assigned to plants in parcels A and B that had developed the first whorl of plagiotropic branches (referred hereafter as jorquette) and possessed mostly branches of >30 cm length. Furthermore, treatments Thin_1 and Thin_2 were only assigned to plants with at least four primary branches, and Control, Thin_1 and Thin_2 treatments were only assigned to plants that had the majority of the apical buds of the primary branches intact.

Treatment allocation, initial baseline measurements, and pruning treatments were carried out at the same time when plants were in the dormant phase of the flushing cycle (i.e., the intermitted production of leaves followed by a period of dormancy; Greathouse et al., 1971). This meant that leaves from previous flushes were fully hardened and terminal buds were not swollen or producing new leaves. As flushing was not synchronized across all plants, treatment allocation and pruning intervention were carried out in multiple batches from September 2020 to December 2020 in parcel A and from November 2020 to January 2021 in parcel B. In each batch, an equal number of plants was assigned to each treatment including control (Table S 1).

### 2.5. Tree architecture description

We measured the diameter of the main stem below the jorquette and the number of primary branches of each selected plant. We then counted the number of internodes (corresponding to the number of axillary buds) and measured the length (cm) of each primary branch of a plant. Finally, we measured the length and noted the position (rank) of all lateral branches on each primary branch, starting from the top of the primary branch. In the Head_66% treatment, we counted and measured secondary branches only on the remaining section of the primary branches, and for Thin_1 and Thin_2 treatments only on the remaining primary branches.

After pruning (or after initial measurement in the case of Control plants) flushing activity was monitored every two weeks to determine when a plant had completed one full flushing cycle. A flush event was considered complete when all the newly formed leaves were fully expanded and dark green in color.

After the completion of one flushing cycle, we conducted the same measurements as described in the previous paragraph. For Control, Thin_1, and Thin_2 plants, we also recorded whether the apical bud of each primary branch was still present (as apical buds can be lost due to insect or physical damage, thus invalidating the treatments).

### 2.6. Data analysis

#### Calculation of variables

Plants from the Control and thinning treatments that had no or damaged apex after one flushing cycle were excluded from the analysis.

Based on the architectural descriptions at the moment of treatment application and after one flushing cycle, we created two binary variables, with values for all internodes: one for the overall presence of lateral branches at the end of the flush and a second variable for branch emergence during the flush (Figure S 2).

We calculated branch length increment of all secondary and primary branches (of Control, Thin_1 and Thin_2) by subtracting the initial branch length from the final branch length.

To calculate total branch-length increment per primary branch we then summed the increment of all secondary branches in a primary branch plus the length increment of the primary branch itself (Control, Thin_1 and Thin_2). Finally, we summed the total increment of all primary branches of each plant to obtain total branch length increment per plant. The last variable was calculated only for those plants for which all primary branches were included in the analysis (i.e., no branch die back and/or no loss of terminal apexes due to e.g. insect damage). Sample sizes for this analysis are given in parenthesis in Table S 1.

#### Statistical analysis

We applied linear generalized mixed effect models (GLMMs, binomial distribution) to explain the variation in lateral branch presence in unpruned plants and lateral branch emergence in all plants. Both branch presence and branch emergence were described as a quadratic function of branch rank, with parcel identity as an additional fixed factor to control for possible effects of differences in local conditions and/or genotype between parcels. For branch emergence we also tested the effect of pruning treatments (Control, Head_tip, Head_66%, Thin_1 and Thin_2), number of lateral branches already present at pruning (N. old branches). All two-way interactions were included in the full model.

To explain the variation of branch length increment and final branch length we applied linear mixed effect models (LMM) as a function of rank and parcel. For lateral branch length increment, we also tested the effect of pruning treatments and their interaction with rank. Finally, branch length increment at primary branch and plant level was tested with LMMs as a function of pruning treatment and parcel only.

In all analyses, we included random intercepts for primary branches nested in plants to account for the nesting of lateral branches on the main branch.

To determine the best-fitting model we tested all possible combinations of fixed effects and their two-way interactions, including an intercept-only model. Model selection was based on Akaike Information Criteria (AIC), which provides an approximation of model predictive accuracy, as measured by out-of-sample deviance (McElreath, 2018). We selected the model with the lowest AIC and in the case multiple models had ΔAIC smaller than 2, the simplest model was selected. Confidence intervals (CI) at 95% of each parameter were calculated as ±1.96 times its standard error. Parameters were considered significant if the CI did not overlap with zero.

If the selected model contained an interaction between pruning treatments and a discrete variable we estimated the slope of the relation for each pruning treatment and tested whether the estimated slopes were significantly different from control.

All analyses were performed using R 4.2.1 (R Core Team 2022). For both GLMMs and LMMs we used the glmmTMB function from the glmmTMB packages. For the post-hoc test we used the emmeans and multcomp packages. To calculate pseudo R^2^ for mixed effect models we use the r.squaredGLMM function from the MuMIn package.

## 3. Results

### 3.1. Branching pattern of unpruned trees

#### Changes in probability of branching along a primary branch

In unpruned trees, branching probability was higher for trees in parcel A compared to parcel B (Figure 2). In both parcels, moving from the tip to the base of a primary branch, branching probability first slightly increased (reaching an estimated maximum of 0.34 at rank 11 for trees in parcel A and 0.17 at rank 33 for trees in parcel B) and then decreased slowly. This pattern was best described by a concave parabola (Figure 2). However, the observed variation explained by the fixed factors was very low.

**Figure 2.**
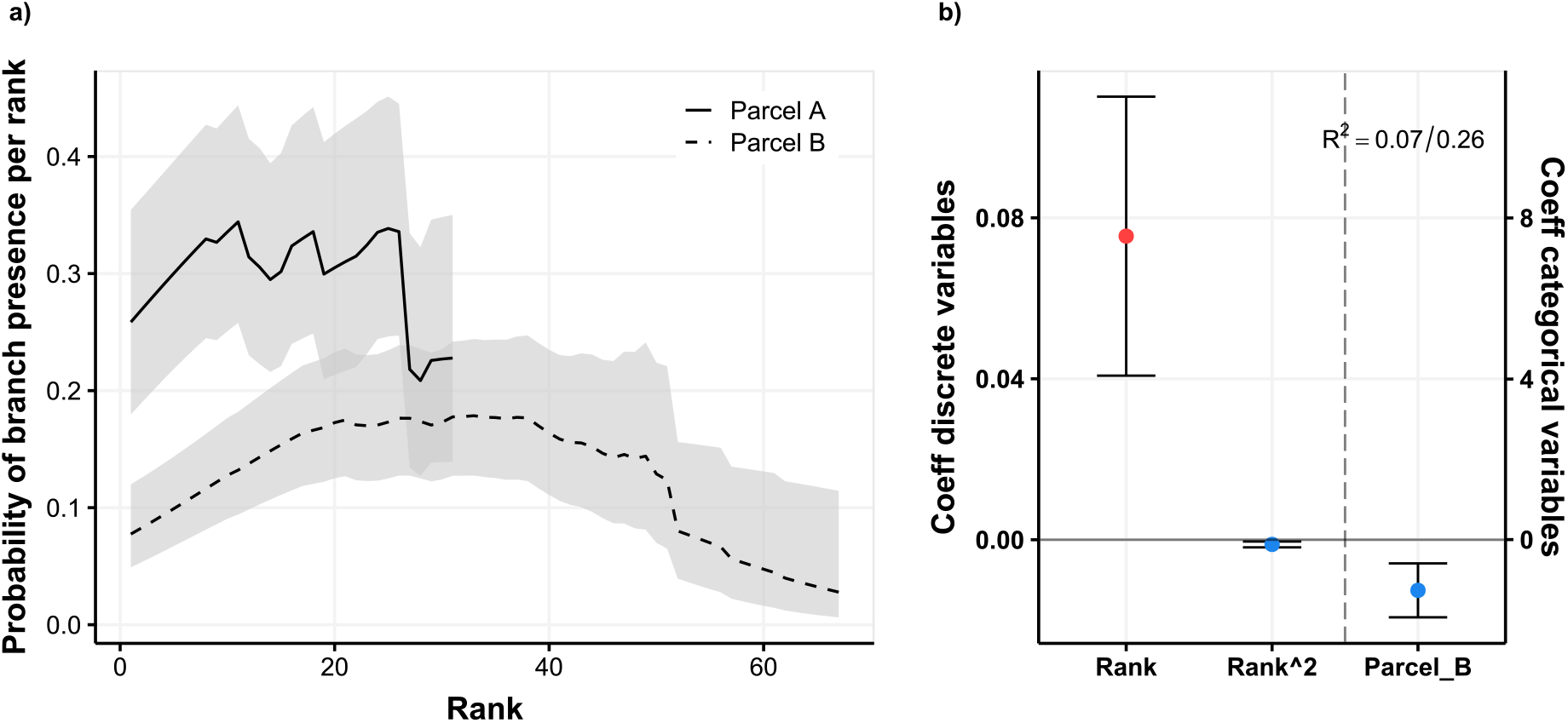
Branching patterns of unpruned cocoa plants. A) Estimated probability of presence of lateral branches along a primary branch in unpruned trees for parcel A (continuous) and parcel B (dashed) using the mixed effect logistic regression. Grey areas indicates 95% confidence interval. B) Coefficients of the mixed effect logistic regression for discrete (left y axis) and categorical (right y axis) variables. Positive significant coefficient shown in red, negative significant coefficient shown in blue. Vertical error bars indicate 95% confidence intervals. Marginal and condition R^2^ are shown.

**Figure 3.**
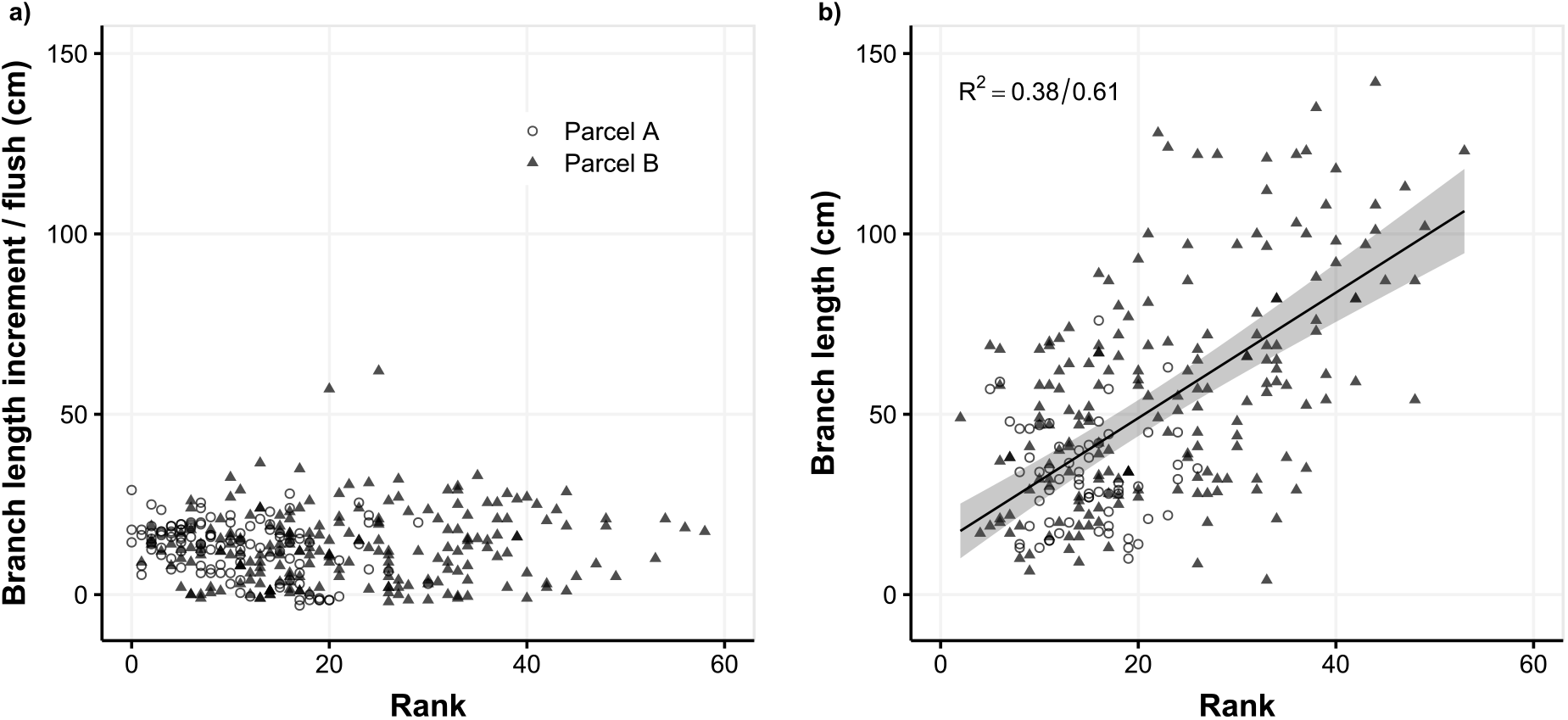
Branch growth and length in unpruned cocoa plants. A) Branch length increase of secondary branches and B) final length of secondary branches. Circles indicate branches of plants from parcel A and triangles plants from parcel B. Model prediction are shown (continuous line indicates a significant relation, dashed line a non-significant relation). 95% confidence interval of model predictions are shown in grey.

#### Lateral branch growth and total branch length along a primary branch

The length increment did not change significantly with rank (slope=0.03, p-value=0.48), implying that during a flushing episode, the length of lateral branches increased at a rate which was unrelated to their position along the primary branch (Figure 4 A).

**Figure 4.**
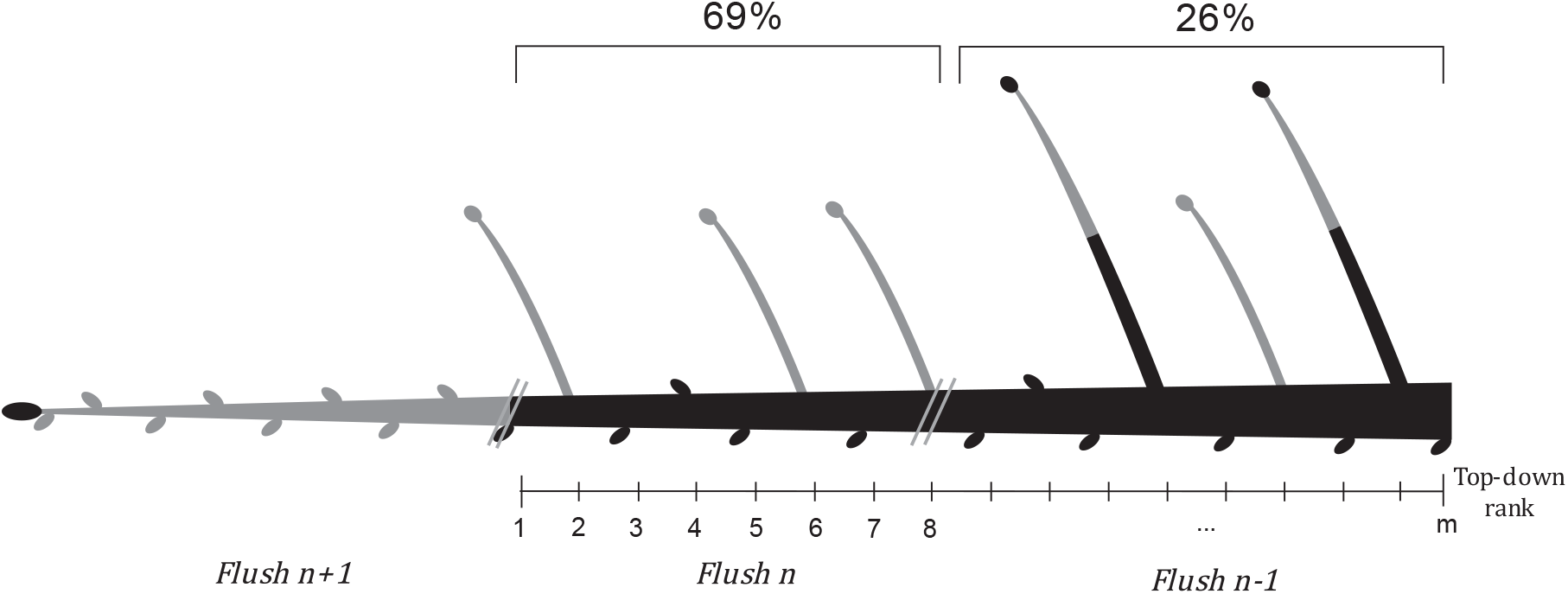
Visualization of emergence of new branches in various sections of a primary branch in unpruned condition. Average percentage of branches emerged in each sections in this study are given. Branch sections produced in the last flush (n+1) are shown in grey.

The total length of lateral branches increased going from the tip to the base of a primary branch (slope=1.71, p-value<0.001; Figure 4 B). This linear model explained a substantial part (38%) of the observed variation. We found no difference in lateral branch length increase and total branch length between the two parcels.

#### 3.2. Emergence of new branches

In unpruned trees, 69% of new lateral branches developed on the first eight internodes of the old branch section (section “flush n” in Figure 4). Those eight internodes correspond roughly to the section of the primary branch produced in the penultimate flushing episode, if we 1) consider that on average eight new internodes were produced per primary branch during the last flush and 2) assume that the number of internodes produced during a flush remains constant during consecutive flushing. A much smaller fraction, 26% of new branches emerged on the older section of the branch (top-down rank > 8). The last 3% was unaccounted for.

#### Pruning effect on branch emergence

Considering all treatments together, our model showed that the probability of branch emergence was higher in the distal part of the branch (excluding section n+1), and quickly decreased moving toward the base of the primary branch. This was best described by a convex parabola (Figure 5a). In addition, we found that the probability of branch emergence decreased with the number of secondary branches already present on the primary branch. The probability of branch emergence was overall lower in parcel B than in parcel A.

**Figure 5.**
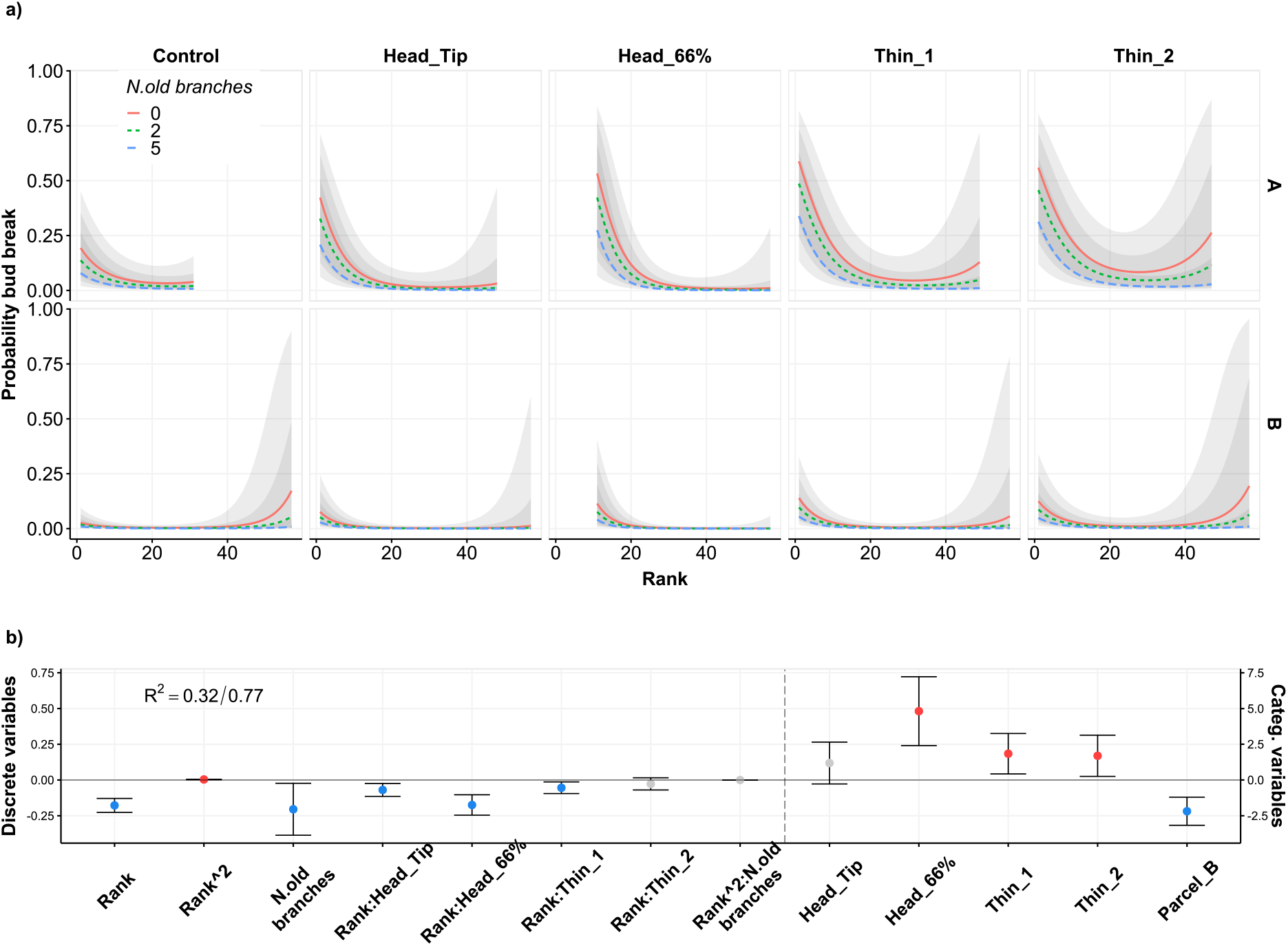
Effect of different pruning treatments on branch emergence in cocoa plants. A) Probability of branch emergence vs branch rank for primary branches with 0 (red continuous line), 2 (green dashed) and 5 (blue dashed) branches, for parcel A (upper panels) and B (lower panels). Grey areas indicates 95% confidence interval. B) Coeffiecients of the mixed effect logististic model for continuous (left axis) and categorical (right axis) variables. Positive significant coefficients shown in red, negative significant coefficient shown in blue, and not significant coefficient in grey. Vertical error bars indicate 95% confidence intervals. ‘Control’ is no pruning, ‘Head_tip’ and ‘Head_66%’ are the removal of the tip and 2/3 of all primary branches, and Thin_1 and Thin_2 are the removal of one or two primary branches, respectively.

All pruning treatments increased the overall probability of branch emergence with Head_66% having the strongest effect (Figure 5b). However, the effect of the removal of the apical bud (Head_Tip) was not significantly different from control (Figure 5b). Except for Thin_2 (the removal of two primary branches), all other pruning treatments had a stronger negative relation with rank than the control (slope=-0.05±0.01), with Head_66% (slope=-0.22±0.03) showing the steepest relation followed by Head_Tip (slope=-0.11±0.01) and Thin_1 (slope=-0.10±0.01) (Tables S 2-S 3). Thus, most pruning treatments concentrated branch emergence more towards the tip of a primary branch compared to control. In Thin_2 instead, the weaker relation with rank and the overall effect on branch probability resulted in an increase of probability of branch emergence also at higher ranks compared to control (Figure 5a).

#### 3.3. Effect of pruning on branch length

In line with the observations on control trees, the length increment of secondary branches after one flushing episode was not influenced by rank and it was also not influenced by pruning treatments. However, contrary to what we observed in control trees, mean branch length increase was larger in parcel A compared to parcel B (A=17.29, B= 11.8, p-value<0.001, Figure 6a).

**Figure 6.**
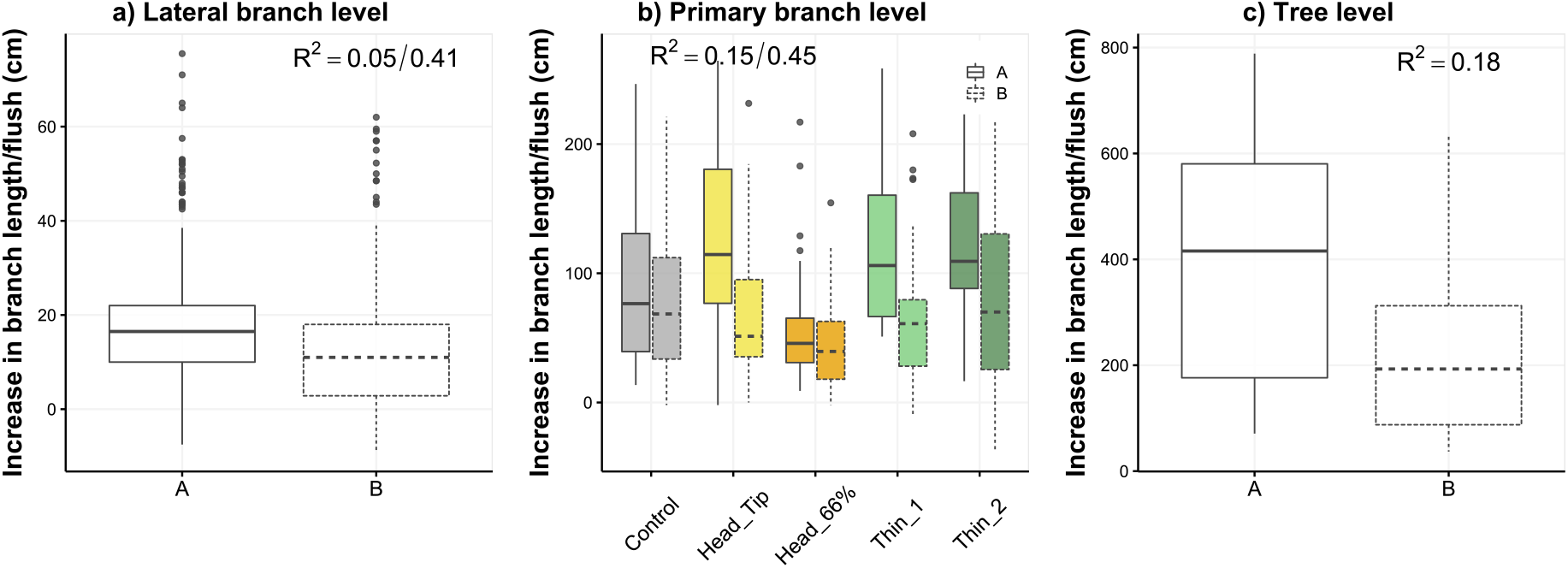
Effects of different pruning treatments on branch growth in cocoa plants. a) Increment in branch length at lateral branch level in parcel A and B. b) Total increment in branch length at primary branch level for each treatment in parcel A (continuous line) and B (dashed line). Colors indicates different treatments. c) Total branch increment at plant level in parcel A and B. Boxplots show the range (rectangle from 25^th^ to 75^th^ percentile), the median (middle line), the calculated minimum (25^th^ percent.-1.5 * the range) and calculated maximum value (75^th^ percent.+ 1.5 * the range, whiskers), and outliers (dots). ‘Control’ is no pruning, ‘Head_tip’ and ‘Head_66%’ are the removal of the tip and 2/3 of all primary branches, and Thin_1 and Thin_2 are the removal of one or two primary branches, respectively.

Head_66% branch length increase at primary branches level was significantly lower than control (Coeff= -34±11.9, p-value<0.01, Figure 6b), while other treatments did not differ significantly from control (Table S 4). The total length increment per primary branch (the sum of the length increments of all secondary branches plus the increment of primary branch length) was significantly higher in parcel A compared to parcel B (A= 104.5, B=71.998, p-values<0.001).

Finally, the total branch length increment per tree was also higher for parcel A compared to parcel B (A= 390.7, B=216.6, p-values<0.001, Figure 6c). The variation explained by the three models was low.

## 4. Discussion

To our knowledge, this study was the first to quantitatively describe branching pattern of plagiotropic branches in cocoa, and to quantitatively assess how pruning intervention changes the pattern of branch emergence. As such it contributes important information needed to develop effective pruning strategies in cocoa and more generally, adds to the knowledge of tree architectural development and its responses to (partial) branch loss.

### 4.1. Cocoa branching pattern

In unpruned trees, the primary plagiotropic branches showed a certain level of lateral branch outgrowth. This can be classified as an “intermediate” level of apical dominance (sensu Cline 1997) meaning that the inhibition imposed by the terminal apex is partial and some axillary buds can grow into a branch when the terminal apex is present.

Considering that cocoa is a shade-tolerant species, we expected a branching pattern that allowed horizontal spreading and minimized leaf overlap, for example with more and longer lateral branches in the distal section of a primary branch (Niinemets, 2010). This expectation was mostly confirmed by our results. In agreement with our expectations, branching probability was lower toward the base of primary branches, the emergence of new branches was concentrated in the distal section of a primary branch, and branching was inhibited if more lateral branches were present. Yet, in contrast to our expectation, we found no preferential allocation to more distal lateral branches, and basal branches were (therefore) longer than more distal branches.

### Formation of the branching pattern

No branches emerged in newly formed sections of a branch. The majority of new branches emerged in the penultimate flush section, a section that at the beginning of the last flushing episode was unbranched. Additionally, fewer branches emerged further away from the growing tip, where some lateral branches were already present. First, this pattern of emergence suggests that axillary buds formed in the last flush need to undergo a short period of dormancy of at least one flushing cycle, before being able to outgrow into a branch. Therefore, lateral branching on the primary horizontally oriented (plagiotropic) branches is to a certain degree proleptic, meaning that the development of the lateral meristems is discontinuous from the terminal meristem (Hallé et al., 1978).

In addition, our results suggest that both within and between branches correlative inhibition plays a role in regulating branching pattern, a pattern that is also observed in roses (Wubs et al., 2013; Zieslin & Halevy, 1976). The emergence of new branches in fact was strongly limited by the presence of other lateral branches. Additionally, the presence of lateral branches decreases slightly at the base of a primary branch, suggesting an inhibitory effect exerted by the other jorquette branches.

Light quality and quantity are also known to influence branching, and buds in different positions of the crown may have experienced different light conditions (Schneider et al., 2019). However, as internal and environmental signaling are deeply interconnected in the regulation of branch outgrowth disentangling those various factors is difficult (McSteen & Leyser, 2005b) and would require dedicated experiments in controlled environment (Wubs et al., 2013), possibly in combination with modelling of the interaction between architecture and light environment (Evers et al 2011).

A possible implication of this branching pattern is that, as the primary branch grows, the lateral branch structure of a section is almost completely determined in the following flushing episode. In later flushes, this structure seems not to change much, except for some occasional branching events. This process creates an age gradient among lateral branches, with older branches at the base and newer branches toward the tip of the branch. Given that we found no difference in lateral branch growth (i.e., rates of length increment) along a primary branch, the branch ages explain the observed gradient in branch length.

Trees in parcel B, all belonging to a single hybrid, showed an overall lower level of branching compared to the trees in parcel A, a mix of several hybrids. Our experimental setup did not allow us to disentangle a possible genotypic effect from a location effect. However, we consider to be unlikely that the location of the parcels would have induced these differences, given that the parcels are less than 150 m from each other and were managed in the same way. Follow-up experiments to characterize the branching pattern of different cocoa hybrids and to test their performance in different shade levels are needed to provide plant material recommendations for different cocoa cropping systems (i.e. full sun vs agroforestry). As branching structure is only one of several factors that determine the light interception efficiency of a tree crown, such experiments should also take into account characteristics such as branch and leaf angle. Additionally, dynamic processes such as branch loss and change in branching pattern with the development of a vertical light gradient in older tree crowns should be included.

### 4.2. Pruning responses to heading and thinning cuts

We hypothesized that heading cuts would induced a strong branching response towards the end of the primary branches, while thinning cut would induced a less strong response, towards the base of the primary branches. The response to pruning varied with pruning treatments and was overall in line with our expectations, except for the fact that thinning cuts induced a stronger response than expected and similar to what observed for the heading cuts.

Heading cuts stimulated the production of new branches in the proximity of the cut. This was more evident when we removed approximately two-third of the branch than when we only removed the terminal apex. Similar results were reported for mango trees, in which the number of lateral branches increased with pruning intensity (Persello et al., 2019). Our results suggest that both apical dominance and within-branch correlative inhibition limit branching, as the removal of the terminal apex alone did not suffice to induce a strong branching response. Yet, our results differ from those in apple trees, where close to 100% of buds in very close proximity of the pruning cut develop into a branch after applying heading cuts (Fumey *et al*., 2011). In our experiment, the probability of a branch emerging on the first rank below the cut did not exeed 53% in parcel A and 11% in parcel B.

Both intensities of thinning cuts stimulated the production of new branches along the entire length of the remaining primary branches. This was more evident when we removed two primary branches than when only one primary branch was removed. The stimulation of branch emergence on the remaining primary branches suggests a release of the between-branches correlative inhibition as also observed in roses (Wubs et al., 2013). The removal of a number of jorquette branches may have also improved the light conditions of the remaining branches and triggered bud break deeper in the crown. Additional experimental and architectural models (Evers et al 2011) are needed to disentangle and quantify those effects.

In our experiment, the two types of pruning cuts did not trigger very contrasting responses. This contradicts with results for other tree crop, such as apple trees, where heading cuts trigger a strong branching response while thinning cuts mostly enhance branch vigor without inducing much branching (Ferree & Schupp, 2003; Fumey et al., 2011). We hypothesize that in cocoa the responses to the two types of pruning cuts may converge as a result of constraints induced by the shade-tolerant nature of the crop. This convergence may result from strong inhibitory signaling between organs. Understory trees need to balance between a modest branching response after the apex loss to avoid leaf overcrowding, on the one hand, while still producing new branches after branch loss to recover total light capture. In tropical forest understory conditions, both leaf overcrowding and a limited capacity of intercepting light reduce growth and survival (Niinemets, 2010).

### 4.3. Lack of compensatory responses after one flushing cycle

As large amounts of biomass were removed in three of the four pruning treatments, we expected plants in these treatments to show a compensatory response (Anten et al., 2003), in terms of an increase in branch growth per flush, a proxy measurement for vegetative production. However we found no clear evidence for such a compensatory response after one flushing episode.

At lateral branch level, the length of a single flush was rather stable and insensitive to perturbances such as pruning. The average number of internodes produced (eight) in a single flush was similar to what we observed in fully developed trees of similar hybrids (A. Tosto, personal observation) and to what was reported by a previous study on cocoa flushing (Greathouse et al., 1971).

Removal of 2/3 of the branches showed a much lower total branch length increase, a possible consequence of strong reduction in total bud number after this intensive pruning.

The lack of a clear compensatory response contrasts with previous research on vegetative responses to pruning in adult cocoa plants, which reported an increase in vegetative production following pruning (Leiva-Rojas et al., 2019; Tosto et al., 2022b). This difference may partially be explained by tree age: adult trees have larger amounts of stored reserves, allowing for more compensatory growth. Longer periods of monitoring biomass and leaf area are likely needed to evaluate evidence for compensatory growth after pruning.

### 4.4. Research outlook and implications

Our study constitutes a first step towards a better understanding of cocoa responses to pruning. As pruning interventions are often a combination of different pruning cuts of different intensities, the interactive effect of various pruning cuts should be assessed in follow-up studies to allow for a more complete understanding of pruning responses at the crown level. In addition, since light availability also plays an important role in branch production and growth (Leduc et al., 2014), pruning responses should be investigated while accounting for the vertical light gradients in mature cocoa crowns and for different shade levels.

Our study provides some practical insights into the manipulation of cocoa crowns. Importantly, we found that in contrast with other fruit trees, pruning responses in cocoa are rather unpredictable. This generates difficulties for practitioners to finely manipulate crown structure through pruning. It also implies that knowledge and pruning strategies developed for other trees may not be directly applicable to cocoa.

For young cocoa trees, our results can be applied to inform formation pruning interventions. For example, in a shaded system, where a strong branching response may results in leaf overcrowding, light heading cuts should be preferred over heavy heading cuts. On the contrary, in full-sun high-density systems, where the goal is to have small compact crowns, heavy heading is preferred. Thinning cuts, that stimulate branching along the entire primary branch, could be used to create open crowns with a more uniform distribution of leaves, which may result in higher light interception efficiency.

Finally, our results can serve as a basis for the development of three-dimensional models of cocoa architecture (Louarn & Song, 2020) that can further our understanding of the performance of cocoa in contrasting shade levels, as well as allowing for exploration of direct and long-term effects of pruning intervention on cocoa functioning.

## 5. Acknowledgments

The authors would like to thank Barry Callebaut for the opportunity to conduct this experiment in the Tiassalé research station. We also would like to thank Kouassi Koffi Brice Herman for his essential work in data collection. We also want to thank Aka Romain, Alexandre Kamisky, Sandrine Emmanuella Gninahophin for their support with field establishment and maintenance. This research was supported by grant W08.250.305 from the Netherlands Foundation for the Advancement of Tropical Research (NWO-WOTRO), Mondelez International and the Wageningen University covid extension grant.

## Notes

### Competing Interest Statement

The authors have declared no competing interest.

## References

Anten, N. P. R., Martínez-Ramos, M., & Ackerly, D. D. (2003). Defoliation and growth in an understory palm: Quantifying the contributions of compensatory responses. Ecology, 84(11), 2905–2918. https://doi.org/10.1890/02-0454

Cline, M. G. (1997). Concepts and terminology of apical dominance. American Journal of Botany, 84(8), 1064–1069. https://doi.org/10.2307/2446149

Ferree, D. C., & Schupp, J. R. (2003). Pruning and Training Physiology. In D. C. Ferree & I. J. Warrington (Eds.), Apples: Botany, Production and Uses (pp. 363–388). CAB international.

Fumey, D., Lauri, P. É., Guédon, Y., Godin, C., & Costes, E. (2011). How young trees cope with removal of whole or parts of shoots: An analysis of local and distant responses to pruning in 1-year-old apple (malus × domestica; rosaceae) trees. American Journal of Botany, 98(11), 1737–1751. https://doi.org/10.3732/ajb.1000231

Greathouse, D. C., Laetsch, W. M., & Phinney, B. O. (1971). The Shoot-growth Rhythm of a Tropical Tree, Theobroma Cacao. American Journal of Botany, 58(4), 281–286.

Hallé, F., Oldeman, R. A. A., & Tomlinson, P. B. (1978). Forests and Vegetation. In Tropical Trees and Forests. https://doi.org/10.1007/978-3-642-81190-6_5

IITA. (2020). Managing Soils for Increased Productivity and Decreased Deforestationin Cocoa. A Training Manual for Field Officers. Version 1.0.

KAU. (1988). First annual report of the Cadbury-KAU Co-operative Cocoa research project, 1987-1988.

KAU. (1989). Second annual report of the Cadbury-KAU Co-operative Cocoa research project.

KAU. (1991). Cadbury-KAU Co-operation research project-Fourth Annual Report 1990-1991. http://fea.kau.edu/announce/STUDENTS.pdf

Lachenaud, P., Sounigo, O., & Sallée, B. (2005). Les cacaoyers spontanés de Guyane française: état des recherches. Acta Botanica Gallica, 152(3), 325–346. https://doi.org/10.1080/12538078.2005.10515493

Leduc, N., Roman, H., Barbier, F., Péron, T., Huché-Thélier, L., Lothier, J., Demotes-Mainard, S., & Sakr, S. (2014). Light signaling in bud outgrowth and branching in plants. In Plants (Vol. 3, Issue 2, pp. 223–250). MDPI AG. https://doi.org/10.3390/plants3020223

Leiva-Rojas, E. I., Gutiérrez-Brito, E. E., Pardo-Macea, C. J., & Ramírez-Pisco, R. (2019). Vegetative and reproductive behavior of cocoa (Theobroma cacao L.) due to pruning | Comportamiento vegetativo y reproductivo del cacao (Theobroma cacao L.) por efecto de la poda. Revista Fitotecnia Mexicana, 42(2), 137–146. https://doi.org/10.35196/rfm.2019.2.137-146

Li, S. H., Zhang, X. P., Meng, Z. Q., & Wang, X. (2010). Responses of peach trees to modified pruning 1. Vegetative growth. New Zealand Journal of Crop and Horticultural Science, 22(4), 401–409. https://doi.org/10.1080/01140671.1994.9513852

Louarn, G., & Song, Y. (2020). Two decades of functional-structural plant modelling: now addressing fundamental questions in systems biology and predictive ecology. Annals of Botany, 126(4), 501–509. https://doi.org/10.1093/aob/mcaa143

Marini, R. P., Specialist, E., & Tech, V. (2020). Pruning Peach Trees. 020.

McElreath, R. (2018). Statistical Rethinking. Statistical Rethinking. https://doi.org/10.1201/9781315372495

McSteen, P., & Leyser, O. (2005a). Shoot branching. Annual Review of Plant Biology, 56, 353–374. https://doi.org/10.1146/annurev.arplant.56.032604.144122

McSteen, P., & Leyser, O. (2005b). Shoot branching. Annual Review of Plant Biology, 56, 353–374. https://doi.org/10.1146/annurev.arplant.56.032604.144122

Mika, A. (1986). Physiological Responses of Fruit Trees to Pruning. In Horticultural Reviews (Vol. 8, pp. 337–378). https://doi.org/10.1002/9781118060810.ch9

Niemenak, N., Cilas, C., Rohsius, C., Bleiholder, H., Meier, U., & Lieberei, R. (2009). Phenological growth stages of cacao plants (Theobroma sp.): Codification and description according to the BBCH scale. Annals of Applied Biology, 156(1), 13–24. https://doi.org/10.1111/j.1744-7348.2009.00356.x

Niinemets, Ü. (2010). A review of light interception in plant stands from leaf to canopy in different plant functional types and in species with varying shade tolerance. Ecological Research, 25(4), 693–714. https://doi.org/10.1007/s11284-010-0712-4

Obeng Adomaa, F., Vellema, S., Slingerland, M., & Asare, R. (2022). The adoption problem is a matter of fit: tracing the travel of pruning practices from research to farm in Ghana’s cocoa sector. Agriculture and Human Values. https://doi.org/10.1007/s10460-021-10292-0

Persello, S., Grechi, I., Boudon, F., & Normand, F. (2019). Nature abhors a vacuum: Deciphering the vegetative reaction of the mango tree to pruning. European Journal of Agronomy, 104, 85–96. https://doi.org/10.1016/j.eja.2019.01.007

Poorter, L., Bongers, L., & Bongers, F. (2006). Architecture of 54 moist-forest tree species: Traits, trade-offs, and functional groups. Ecology, 87(5), 1289–1301. https://doi.org/10.1890/0012-9658(2006)87[1289:AOMTST]2.0.CO;2

Sachs, T., & Novoplansky, A. (1995). Tree form: Architectural Models do Not Suffice. Israel Journal of Plant Sciences, 43(3), 203–212. https://doi.org/10.1080/07929978.1995.10676605

Schneider, A., Godin, C., Boudon, F., Demotes-Mainard, S., Sakr, S., & Bertheloot, J. (2019). Light Regulation of Axillary Bud Outgrowth Along Plant Axes: An Overview of the Roles of Sugars and Hormones. Frontiers in Plant Science, 10. https://doi.org/10.3389/fpls.2019.01296

Tosto, A., Zuidema, P. A., Goudsmit, E., Evers, J. B., & Anten, N. P. R. (2022a). The effect of pruning on yield of cocoa trees is mediated by tree size and tree competition. Scientia Horticulturae, 304(October 2021), 111275. https://doi.org/10.1016/j.scienta.2022.111275

Tosto, A., Zuidema, P. A., Goudsmit, E., Evers, J. B., & Anten, N. P. R. (2022b). The effect of pruning on yield of cocoa trees is mediated by tree size and tree competition. Scientia Horticulturae, 304(October 2021), 111275. https://doi.org/10.1016/j.scienta.2022.111275

Wilson, B. F. (2000). Apical control of branch growth and angle in woody plants. American Journal of Botany, 87(5), 601–607. https://doi.org/10.2307/2656846

Wubs, A. M., Heuvelink, E., Marcelis, L. F. M., Okello, R. C. O., Shlyuykova, A., Buck-Sorlin, G. H., & Vos, J. (2013). Four Hypotheses to Explain Axillary Budbreak after Removal of Flower Shoots in a Cut-rose Crop. J. Amer. Soc. Hort. Sci., 138(4), 243–252. http://journal.ashspublications.org/content/138/4/243.abstract#fn-4

Zieslin, N., & Halevy, A. H. (1976). Components of Axillary Bud Inhibition in Rose Plants. I. The Effect of Different Plant Parts (Correlative Inhibition). Botanical Gazette, 137(4), 291–296.

