## Supplemental methods info and tables for "Branching responses to pruning in cocoa"

### Supplementary material

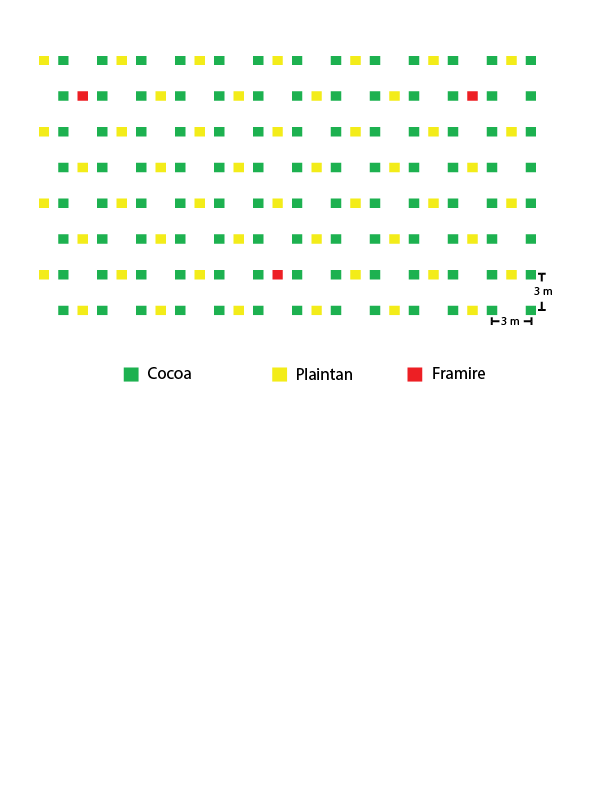

Figure S 1 Planting design of parcel A and parcel B.

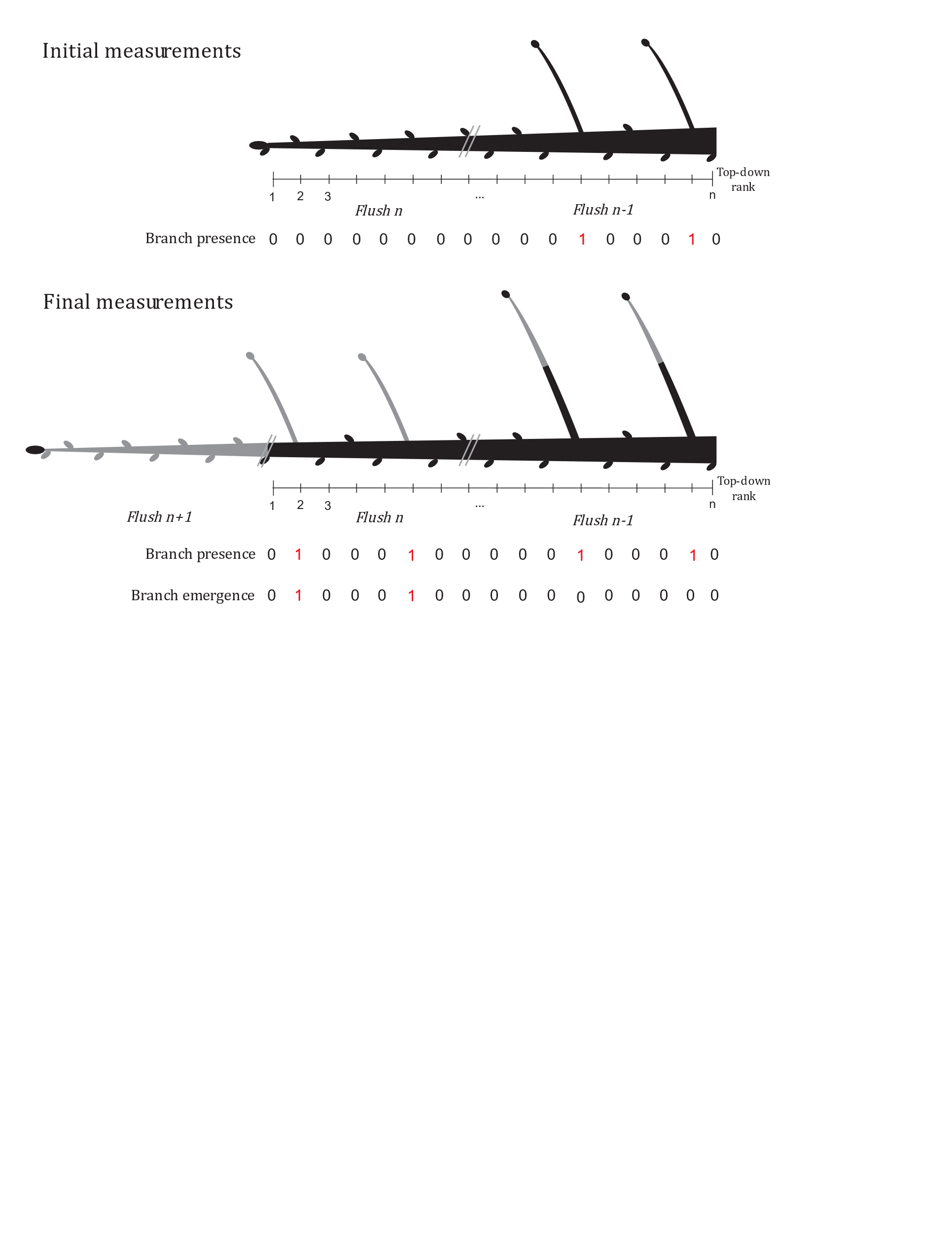

Figure S 2 Visual representation of primary branches, with example of binary variables for branch presence and branch emergence. Branch sections produced in the last flush (n+1) are shown in grey.

Table S 1 Overview of number of plants and number of primary branches per treatment for the two parcels. In parentesis are number of plants for which all branches were used in the analysis, this is also the sample size for the plant-level analysis of branch length increase.

|  | **Number of plants** | | | **Number of primary branches** | | |
| --- | --- | --- | --- | --- | --- | --- |
|  | **Parcel A** | **Parcel B** | **Total** | **Parcel A** | **Parcel B** | **Total** |
| **Control** | 7 (4) | 15 (8) | 22 (12) | 22 | 43 | 65 |
| **Head_Tip** | 6 | 9 (8) | 15 (14) | 23 | 39 | 62 |
| **Head_66%** | 10 (6) | 16 (14) | 26 (20) | 40 | 58 | 98 |
| **Thin_1** | 6 (2) | 17 (8) | 23 (10) | 18 | 52 | 70 |
| **Thin_2** | 9 (4) | 15 (9) | 24 (13) | 22 | 33 | 55 |

Table S 2 Estimated slope and standard error of effect of rank on probability of branch emergence per pruning treatments.

| **Treatments** | **Estimated effect of Rank** |
| --- | --- |
| Control | -0.05±0.01 |
| Head_Tip | -0.12±0.01 |
| Head_66% | -0.22±0.03 |
| Thin_1 | -0.10±0.01 |
| Thin_2 | -0.08±0.02 |

Table S 3 Control versus treatment tests for the effect of rank on probability of branch emergence.

| **Treatment-contrast** | **Difference estimates** | **t- ratio** | **df** | **p.value** |
| --- | --- | --- | --- | --- |
| Head_Tip - Control | -0.07±0.02 | -3.01 | 9016 | 0.009 |
| Head_66% - Control | -0.17±0.03 | -4.805 | 9016 | <0.001 |
| Thin_1 - Control | -0.05±0.02 | -2.61 | 9016 | 0.033 |
| Thin_2 - Control | -0.02±0.02 | -1.2 | 9016 | 0.52 |
| P value adjusted with Dunnettx methods for 4 tests. | | | | |

Table S 4 Results of mixed effect models for branch length increase at primary branches level. Shown are mean and standard error (SE) of model coefficients, Z values and p.values.

| **Treatment** | **Coeff. Estimates** | **Z value** | **p.value** |
| --- | --- | --- | --- |
| Control-Par A (Intercept) | 104.5±10.6 | 9.844 | <0.001 |
| Head_Tip | 1.8±13.5 | 0.130 | 0.896 |
| Head_66% | -34.6±11.9 | -2.898 | 0.003 |
| Thin_1 | -4.5±12.4 | -0.360 | 0.718 |
| Thin2 | 14.4±12.7 | 1.132 | 0.257 |
| Parcel B | -32.5±8.3 | -3.901 | <0.001 |
